# Systematic analysis of genetic and phenotypic characteristics reveals antisense oligonucleotide therapy potential for one-third of neurodevelopmental disorders

**DOI:** 10.1101/2025.03.20.644369

**Authors:** Kim N. Wijnant, Nael Nadif Kasri, Lisenka ELM Vissers

**Affiliations:** Department of Human Genetics, Radboud University Medical Center, Nijmegen, The Netherlands; Donders Institute for Brain, Cognition and Behaviour, Radboud University Medical Center, Nijmegen, The Netherlands; Research Institute for Medical Innovation, Radboud University Medical Center, Nijmegen, the Netherlands

**Keywords:** Antisense oligonucleotides, Neurodevelopmental disorders, AON therapy suitability, Targetability assessment

## Abstract

**Background:** Neurodevelopmental disorders (NDDs) are a challenging group of disorders to treat, but promising therapeutic interventions in the form of antisense oligonucleotides (AONs) have emerged in recent years. However, the applicability of AON therapy for NDDs varies based on genetic and phenotypic traits. In this study we systematically evaluated key characteristics for AON therapy suitability in NDDs, to estimate overall therapy potential and identify, both well- and less-studied, targetable NDDs.

**Methods:** An NDD dataset was created and evaluated to identify potentially targetable NDDs for seven AON strategies. This involved examining the presence of a combination of critical factors including disease-gene properties, such as regulatory elements, effects of pathogenic variants, and disease-associated phenotypic features.

**Results:** Through systematic evaluation of the presence of targetable characteristic for each NDD and AON strategy, we identified 711 NDDs (38% of the total) with characteristics favorable for at least one AON strategy and predicted that 18% of affected individuals could benefit from AON therapy.

**Conclusion:** The results from our analysis demonstrate that there might be a more extensive potential for the use of AON therapy in NDDs than was anticipated thus far, underscoring AON therapy as a promising treatment option for NDDs while simultaneously contributing to informed therapy selection.

## Background

Treatment of neurodevelopmental disorders (NDDs) has been exceptionally challenging. Many NDDs manifest early during development and it remains to be elucidated which clinical features associated with NDDs, such as structural abnormalities in the brain, can be reversed in children and adults after manifestation(1). In addition, various aspects of NDDs further complicate therapy development, including the delivery over the blood-brain barrier, the vast array of NDDs - each with its individually rare occurrence, diverse, complex, and often poorly understood mechanism of disease - as well as the phenotypic and genetic heterogeneity. In the last decade, however, antisense oligonucleotide (AON)-based therapies have emerged, also as promising therapy for NDDs(2).

AONs are short oligonucleotides that bind the (pre-)mRNA and can be used to affect splicing, stability, and translation (Fig. 1). Splice-switching AONs can (partially) restore a mutated gene’s function by rescuing splice-disrupting variants(3, 4) (Fig. 1A, Supplementary Note 1) or exon-skipping to prevent nonsense-mediated decay (NMD) and allow expression of a shortened transcript by skipping exons containing truncating variants (exon-skipping for variant exclusion)(5) or by skipping out-of-frame exons to restore the reading frame of out-of-frame deletions (exon skipping for reading frame restoration)(6, 7) (Fig. 1B/C, Supplementary Note 1). Furthermore, steric-hindrance AONs can also be used to improve the translation efficiency of the wild-type allele in diseases caused by haploinsufficiency or imprinting by affecting stability or translation efficiency by binding to upstream AUGs, stem structures in the 5’UTR, AU-rich elements in the 3’UTR(8-11), or by targeting NMD-sensitive events to upregulate expression (targeted augmentation of nuclear gene output, TANGO)(12, 13) (Fig. 1D-F, Supplementary Note 1). Alternatively, improved translation efficiency of the wild-type allele can also be achieved by RNAseH1 activating AONs by promoting the degradation of regulatory factors, such as cis-acting long non-coding (lnc)RNAs(14, 15) (Fig. 1G, Supplementary Note 1). Lastly, RNAseH1-activating AONs can also be used to degrade overexpressed, overactive or toxic gene products(16-22) (Fig. 1H). Since AONs bind cellular RNA via complementary base pairing, it is a highly programmable approach to target a pathway, disease, or a specific variant at the root of the problem. Additionally, intrathecal or intracerebroventricular administration of AONs allows central nervous system targeting without crossing the blood-brain barrier(2). These characteristics combined make AONs a promising treatment option for neurological disabilities, including NDDs.

**Fig. 1.**
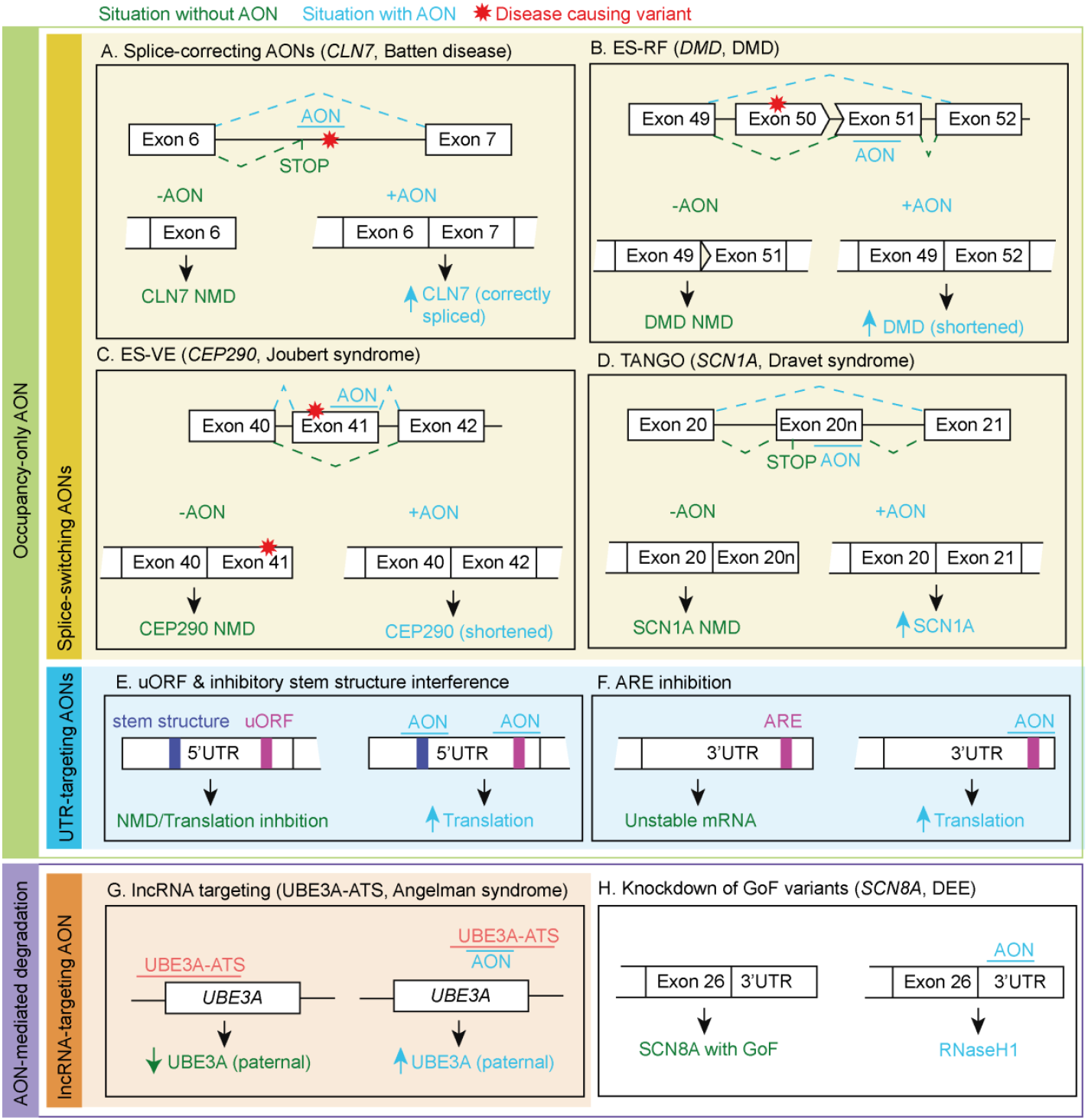
Mechanisms of AON strategies in neurological disorders. A. Batten’s disease, the AON rescues aberrant splicing of intron 6 by steric hindrance of a cryptic splice-acceptor site(3). B. Duchenne muscular dystrophy (DMD), the AON modulates exon skipping of exon 51 directly upstream of a deleted exon 50 for reading frame restoration(5). C. Joubert syndrome, the AON mediates skipping of an exon harboring a truncating variant(7). D. Dravet syndrome, the AON prevents the production of the non-productive splice product containing the poison exon 20N and thereby upregulates expression(12). E. The AON blocks stem structures or uAUGs in the 5’UTR to increase translation efficiency(10). F. The AON blocks an ARE in the 3’UTR to increase translation efficiency(11). G. Angelman syndrome, the AON blocks the transcription of the lncRNA UBE3A-ATS and increases paternal *UBE3A* by inhibiting transcriptional interference(50). H. *SCN8A* DEE, the AON causes RNaseH1-mediated degradation of *SCN8A* mRNA(16). NMD: Nonsense-mediated decay, ARE: AU-rich elements, ES-VE: exon skipping for variant exclusion, ES-RF: exon skipping for reading frame restoration, uAUG: upstream start codon.

Although AON therapy for NDDs is a fast-growing field of interest, not all genetic variants and NDDs are equally suitable for AON-based therapy and it is crucial to consider factors, such as clinical phenotype, causative variant type and certain specific gene properties(23). In terms of clinical phenotype, progressive neurologic phenotypes and seizures are often viewed as an opportunity for therapeutic intervention because they involve ongoing dynamic processes that can be influenced or stabilized with treatment. In contrast, brain malformations present at birth represent fixed structural abnormalities that typically cannot be modified through therapeutic approaches. The emergence of this concept is evident from the AON-based drugs for neurological diseases currently approved and registered by the FDA and EMA, which have a predominant focus on neurodegenerative diseases and DEE-type disorders(24-28). Regarding the type of variant, some splice-disrupting variants can be corrected through splice-switching AONs, while NDDs resulting from heterozygous loss-of-function (LoF) variants in haploinsufficient genes may be targeted using methods to enhance the gene output of the wild-type allele, such as TANGO, or by targeting inhibitory elements in the UTRs or lncRNAs(10, 12, 15, 29). Furthermore, the effectiveness of the latter group of strategies hinges on specific gene properties, such as the presence of NMD-sensitive events for TANGO, inhibitory elements in UTRs or inhibitory lncRNAs. Due to these contributing factors, it is not yet clear what the actual impact of these different AON strategies on the treatment of NDDs can be.

We here set out for a systematic approach to assess which NDDs can be targeted by an AON strategy. Hereto, we created a comprehensive overview of all NDDs and their reported pathogenic variants. Based on the molecular and clinical characteristics of the NDDs, we subsequently determined whether these can be targeted by one or more AON strategies to gain insight into the AON-based therapeutic potential of NDDs.

## Materials and methods

### Creation of the NDD dataset

For the creation of our dataset of NDDs that are amenable for AON therapy, we included entity-level data from the European Reference Network for Rare Malformation Syndromes, Intellectual and Other Neurodevelopmental Disorders (ERN-ITHACA; sysNDD), the Developmental Disorders Genotype-to-Phenotype (DDG2P) database from the UK Deciphering Developmental Disorders project (DDD), the Human Phenotype Ontology (HPO; HPOmim) and our in-house diagnostic gene-panel entries used for ID and epilepsy (Genome Diagnostics Nijmegen)(30-32) (Table 1). For the DDD; DDG2P; entries with Brain/Cognition listed as their affected organ system were selected. We combined and deduplicated the data in the above-mentioned sources on OMIM Disease ID, HGNC ID and inheritance if known and manually curated a selection of NDDs that could not be deduplicated computationally. Each disease-gene association was kept separate, and the clinical validity of the disease-gene associations was ranked based on the cumulative presence of the multiple resource(s) (Supplementary Figure 1)(33). A score of +1 was given to sysNDD entries with category limited and moderate, all HPO entries, DDG2P entries with confidence category limited, moderate, strong, both RD and IF and a score of +2 was given to entries from sysNDD and DDG2P with confidence category definitive and entries from the Genome Diagnostics Nijmegen Exome Panel(33). To ensure the inclusion of only true NDDs into our analysis as much as possible we only considered disease-gene associations with a source-score of 3 and higher. Finally, we excluded 111 entries from our impact analysis that did not have a linked OMIM Disease ID, needed for further annotation.

**Table 1.**
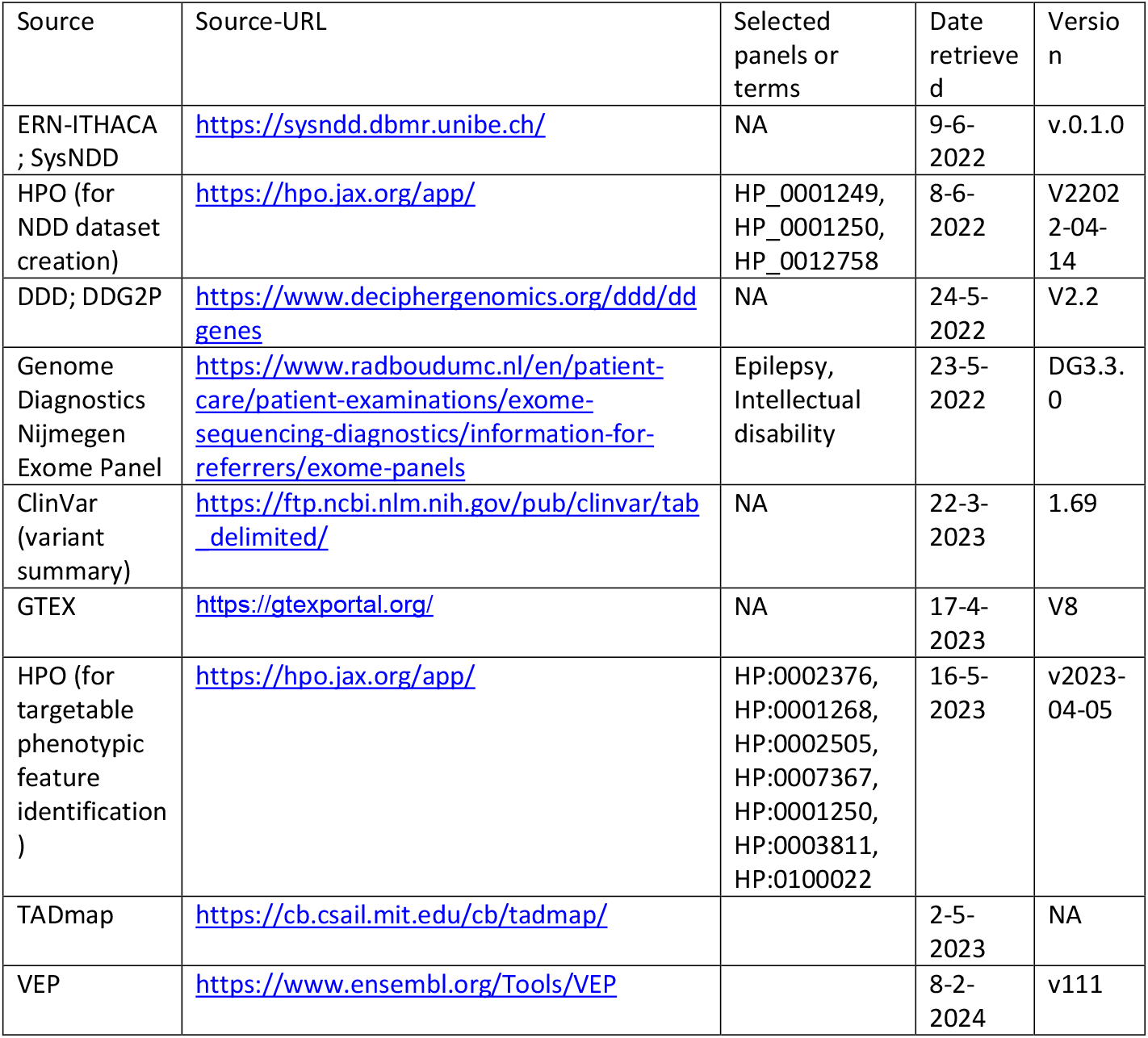
Online resources.

| Source | Source-URL | Selected panels or terms | Date retrieved | Version |
| --- | --- | --- | --- | --- |
| ERN-ITHACA ; SysNDD | https://sysndd.dbmr.unibe.ch/ | NA | 9-6-2022 | v.0.1.0 |
| HPO (for NDD dataset creation) | https://hpo.jax.org/app/ | HP_0001249, HP_0001250, HP_0012758 | 8-6-2022 | V22022-04-14 |
| DDD; DDG2P | https://www.deciphergenomics.org/ddd/ddgenes | NA | 24-5-2022 | V2.2 |
| Genome Diagnostics Nijmegen Exome Panel | https://www.radboudumc.nl/en/patient-care/patient-examinations/exome-sequencing-diagnostics/information-for-referrers/exome-panels | Epilepsy, Intellectual disability | 23-5-2022 | DG3.3.0 |
| ClinVar (variant summary) | https://ftp.ncbi.nlm.nih.gov/pub/clinvar/tab_delimited/ | NA | 22-3-2023 | 1.69 |
| GTEX | https://gtexportal.org/ | NA | 17-4-2023 | V8 |
| HPO (for targetable phenotypic feature identification ) | https://hpo.jax.org/app/ | HP:0002376, HP:0001268, HP:0002505, HP:0007367, HP:0001250, HP:0003811, HP:0100022 | 16-5-2023 | v2023-04-05 |
| TADmap | https://cb.csail.mit.edu/cb/tadmap/ |  | 2-5-2023 | NA |
| VEP | https://www.ensembl.org/Tools/VEP |  | 8-2-2024 | v111 |

### Annotation of ClinVar Variants

We intersected our NDD dataset with ClinVar variants in a single gene with a ClinSigSimple of 1 from ClinVar(34), based on the most reported OMIM disease ID(s) for each variant (Table 1). Ensembl’s variant effect prediction (VEP) with NMD and SpliceAI plugin was used to annotate all ClinVar variants (Table 1). Ambiguous VEP consequences in the splice region were annotated as splice_region_variants if any SpliceAI score > 0.2. Additionally, due to limitations of variant effect prediction for variants of 50 base pairs and longer, these variants were annotated as SVs. For ambiguous VEP consequences in the splice region with a spliceAI score of <0.2 and all other ambiguous VEP consequences the most severe consequence was used to determine the variant type.

### Identification of targetable phenotypic features

To identify NDDs with phenotypic features that are potentially rescuable for AON therapy we annotated our NDD dataset with HPO phenotype terms based on OMIM disease ID and selected all NDDs in our dataset associated with developmental, mental or ambulation deterioration (HP:0002376, HP:0001268, HP:0002505), neurodegeneration (HP:0007367), seizures (HP:0001250) or abnormality of movement (HP:0100022) and all higher-order HPO terms that fall underneath these HPO terms (Table 1, Supplementary Data 3). Subsequently, we excluded all NDDs associated with neonatal lethality (HP:0003811) from the list of NDDs with a likely targetable phenotypic feature. The UpSet plot visualizing the contribution of each HPO-term to the total number of NDDs with a targetable phenotype was created using the ComplexUpset R package with mode distinct(35).

### Identification of targetable variants for splice-correction

To identify deep intronic variants amenable to AON-mediated splice correction we selected all variants with the predicted consequence ‘intron_variant’ located at a distance of 50 bps or more from an exon-intron boundary in the MANE transcript. Subsequently, to identify recurrent variants we selected all amenable deep intronic variants with 3 or more ClinVar submissions.

### Identification of targetable variants for exon skipping

To identify targetable variants for exon skipping to restore the reading frame, we focused on ClinVar variants of the SV type that cause a complete deletion of one or multiple exons, since smaller frameshifts may introduce premature termination codons (PTCs) within the variant-containing exon. In this latter scenario, skipping the adjacent downstream (out-of-frame) exon would not prevent the formation of the PTC (and trigger NMD), and skipping of an adjacent upstream out-of-frame exon would cause a frame-shift in the start of the affected exon, also leading to a PTC and subsequent NMD. Additionally, we selected splice variants with the predicted consequence ‘splice_donor_variant’ or ‘splice_acceptor_variant’ that were located next to an out-of-frame exon, essentially achieving the same effect as the deletion of a full exon. Next, we selected all exons neighboring the deleted exon(s) that had a length that is not a multiple of three but when combined with the length of the deleted exon(s) and the adjacent out-of-frame exon, resulting in a total length that was a multiple of three, therefore resulting in a larger but in-frame deletion according to the principles of exon skipping for reading frame restoration.

To identify variants targetable for exon skipping for variant exclusion we selected ClinVar variants with consequence ‘frameshift_variant’ or ‘stop_gained’ in a single in-frame exon. Although missense variants can, in theory, also be targeted using this approach, we excluded this variant type as the presence of a pathogenic missense variant typically implies a critical role of the respective exon for protein function, making them less suitable for exon-skipping for variant exclusion(36).

For the two exon skipping strategies described above we excluded all variants that result in a deletion of the first and last exon for biological reasons and selected only variants that lead to a maximum of a 10% loss in coding sequence (both the variant, if applicable, and skipped exon combined), based on the MANE transcript and protein.

### Identification of targetable gene properties for wildtype allele upregulation

For the identification of targetable variants with a heterozygous LoF mechanism, we selected LoF variants annotated to an NDD with AD inheritance and located in a gene with a pLI score of > 0.9 where more than 20% of the causative variants led to LoF (37, 38). LoF variants were defined as variants with the predicted consequences of stop-gained, frame-shift or splice donor/acceptor variants, excluding the variants that are predicted to escape NMD. To determine the possibilities for TANGO and uAUG- and ARE-targeting AONs, we intersected a set of 7,758 genes for which non-productive splice products were previously reported (12) and a dataset of 7,826 translated uAUGs in the 5’UTR and overlapping the coding regions in 4,396 genes and all 3’UTR entries in 4,484 genes in ARED-plus(39, 40) with our NDD dataset. For the identification of regulatory cis-acting lncRNAs we determined for all NDD genes whether lncRNAs (based on the biotype as reported in Ensembl [retrieved: 3-5-2023]) are located within the same topologically associated domain (TAD) as defined in the TADmap(41, 42) (Table 1). Subsequently, we assessed the co-expression of these NDD genes and all lncRNAs (with at least one tissue with TPM > 0) within its TAD using brain expression data cataloged in GTEX median gene-level TPM bulk tissue expression data(43) (Table 1). For statistical testing, we performed a Pearson correlation test for each lncRNA-gene pair and corrected for multiple testing using the Benjamini-Hochberg procedure.

### Integration of targetable characteristics

To identify the number of amenable NDDs for different AON strategies we integrated the evaluated gene properties, causative variant types and phenotypes using the ComplexUpset R package with mode intersect(35). An NDD was considered targetable if at least one variant annotated to the respective NDD had a combination of targetable characteristics for at least one AON therapy. To estimate the number of individuals we took the number of submissions in ClinVar per variant as a measure of occurrence. We again integrated the evaluated gene properties, causative variant types and phenotypes using the ComplexUpset package with mode intersect(35). A variant was considered targetable if it had a targetable variant type and the disease and gene annotated to it contained a combination of targetable characteristics for the respective AON strategy. For the total number of targetable NDDs and individuals, all NDDs or individuals with at least one potential AON strategy were taken.

### Calculation of number of amenable individuals per AON

To estimate the average number of individuals per AON given a specific AON strategy, we assessed the total number of amenable individuals based on variant submissions. A variant was deemed amenable if its variant type, gene properties, and phenotype were targetable. The number of AONs needed to target all amenable variants for a specific AON strategy was calculated differently for each strategy:

- Splice-correction (of recurrent variants): for AON-mediated splice-correction, we used the count of unique targetable variants as a measure of the predicted AONs needed to address all amenable individuals.
- Exon-skipping for reading frame restoration and variant exclusion: For exon-skipping for reading frame restoration and variant exclusion, we used the count of all unique exons to skip per gene to target all variants.
- TANGO/uAUG-interference/ARE inhibition/lncRNA targeting: For TANGO/uAUG- interference/ARE inhibition/lncRNA targeting we considered the count of different genes harboring variants relevant to each specific strategy.

## Results

### Creation of the NDD dataset

An overarching dataset of NDDs with relevant information on phenotype, gene properties and causative variants is essential to facilitate a bottom-up assessment of the potential of AON strategies for NDDs. While several databases cataloging information on NDDs exist(30, 31), they are not suited for this purpose as they either do not take allelic disorders into account, lack detailed data at variant level concerning the genetic cause of a disease or do not report on phenotypic traits. To address this gap, we first created a comprehensive dataset of NDD-gene associations with relevant information on phenotype, gene properties and associated variants.

### Defining disease-gene associations

To create a dataset of NDD-gene associations with relevant information on phenotype, gene properties and associated variants, we combined NDD entries obtained from the European Reference Network for Rare Malformation Syndromes, Intellectual and Other Neurodevelopmental Disorders (ERN-ITHACA; sysNDD), the Developmental Disorders Genotype-to-Phenotype (DDG2P) database from the UK Deciphering Developmental Disorders project (DDD), The Human Phenotype Ontology (HPO; HPOmim) and our in-house diagnostic gene-panel entries used for ID and epilepsy (Genome Diagnostics Nijmegen)(30-32). This resulted in a dataset of 1,996 gene-NDD associations in 1,773 unique genes while retaining reported disease features, such as inheritance and OMIM Disease ID (Fig. 2A, Supplementary Data 1). Among the 1,996 gene-disease annotations, 111 lacked a linked OMIM Disease ID, preventing annotation with causative ClinVar variants or HPO phenotype terms. Consequently, these NDDs were excluded from further calculations, resulting in a final dataset of 1,885 disease-gene associations.

**Fig. 2.**
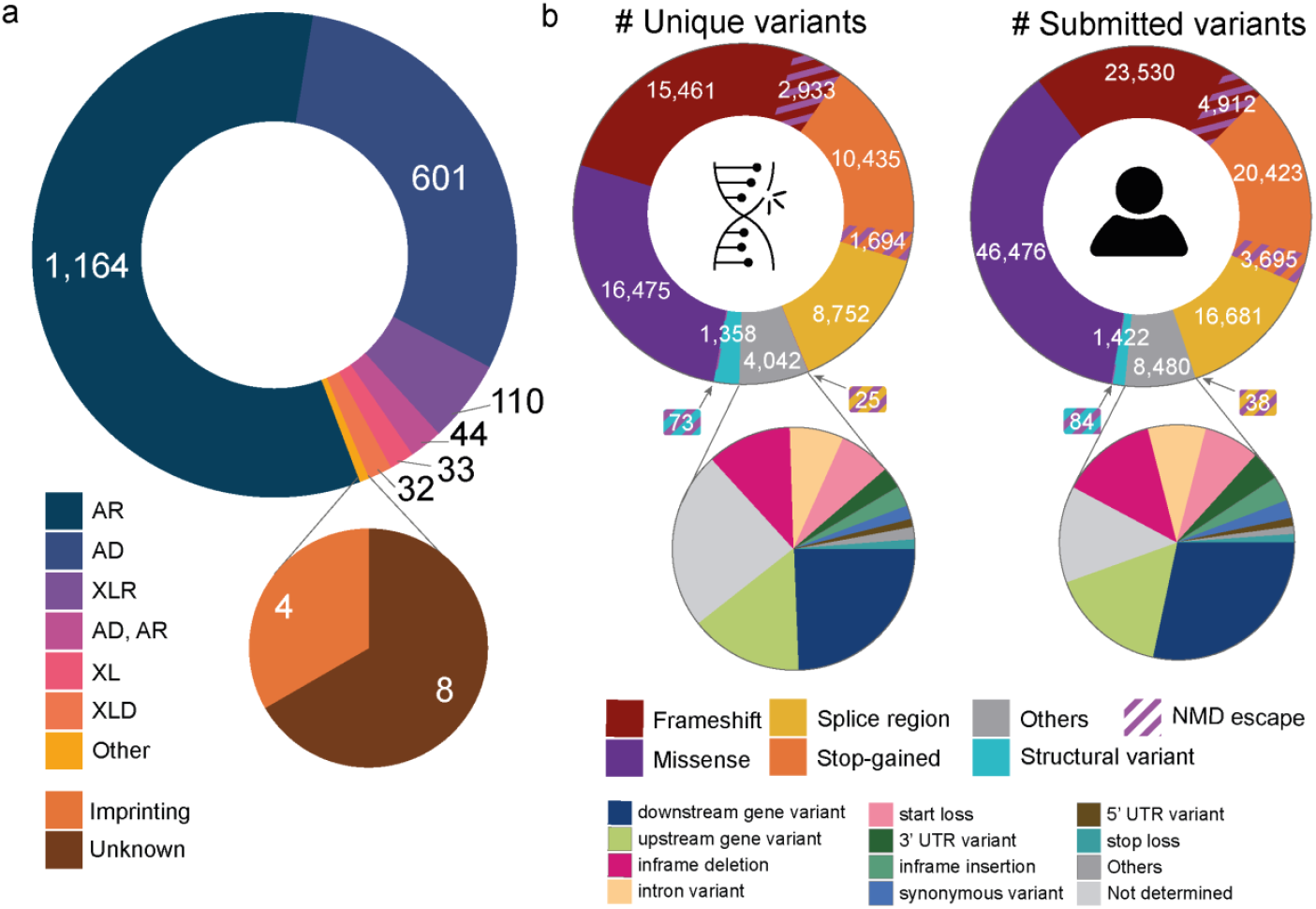
Graphical overview of the NDD dataset. A. The reported inheritance of NDD disease-gene associations B. The predicted variant consequence of all unique ClinVar variants and submissions. AR: autosomal recessive, AD: autosomal dominant, XLR: X-linked recessive, XL: X-linked recessive or dominant, XLD: X-linked dominant, NMD: nonsense-mediated decay, UTR: untranslated region.

### Annotation and classification of ClinVar variants

To assess the variant types associated with the entries in our NDD dataset, we annotated all NDDs in our dataset with related (likely) pathogenic ClinVar variants using OMIM Disease IDs reported in the source databases. This resulted in 61,247 ClinVar variants (125,741 ClinVar submissions) (Fig. 2B). Subsequently, we predicted the consequence of all ClinVar variants based on the MANE transcript and protein using ensemble’s variant effect predictor (VEP)(44, 45) (Fig. 2B). For 306 ClinVar variants (451 submissions) the variant consequence could not be determined and were therefore excluded from further analysis, resulting in a final dataset of 60,941 ClinVar variants with 125,290 submissions (Supplementary Data 2).

### Annotation and identification of targetable phenotypic presentations

To assess the targetability of phenotypic presentations of NDDs, we annotated all NDDs in our dataset with HPO phenotypic terms that are presumed to impact NDD targetability using OMIM Disease IDs reported in the source databases. This involved the incorporation of HPO phenotypic terms for progressive phenotypes and seizures while excluding NDDs associated with HPO phenotypic terms for neonatal lethality (Supplementary Data 3). This led to the identification of 1,408 unique disease-gene associations (75%) with a presumable targetable phenotypic feature (Fig. 3). Of note, as a proof-of-concept of this approach, we noted that these included six NDDs for which clinical trials have been completed or are ongoing like Duchenne Muscular Dystrophy (DMD; OMIM: 310200), Spinal Muscular Dystrophy (SMA; OMIM: 253300), Dravet syndrome (DS; OMIM: 607208), Angelman syndrome (AS; OMIM: 105830), Batten disease (*CLN7*-type, OMIM: 610951) and Ataxia-telangiectasia (AT; OMIM: 208900)(3, 24-28, 46) (Supplementary Data 1).

**Fig. 3.**
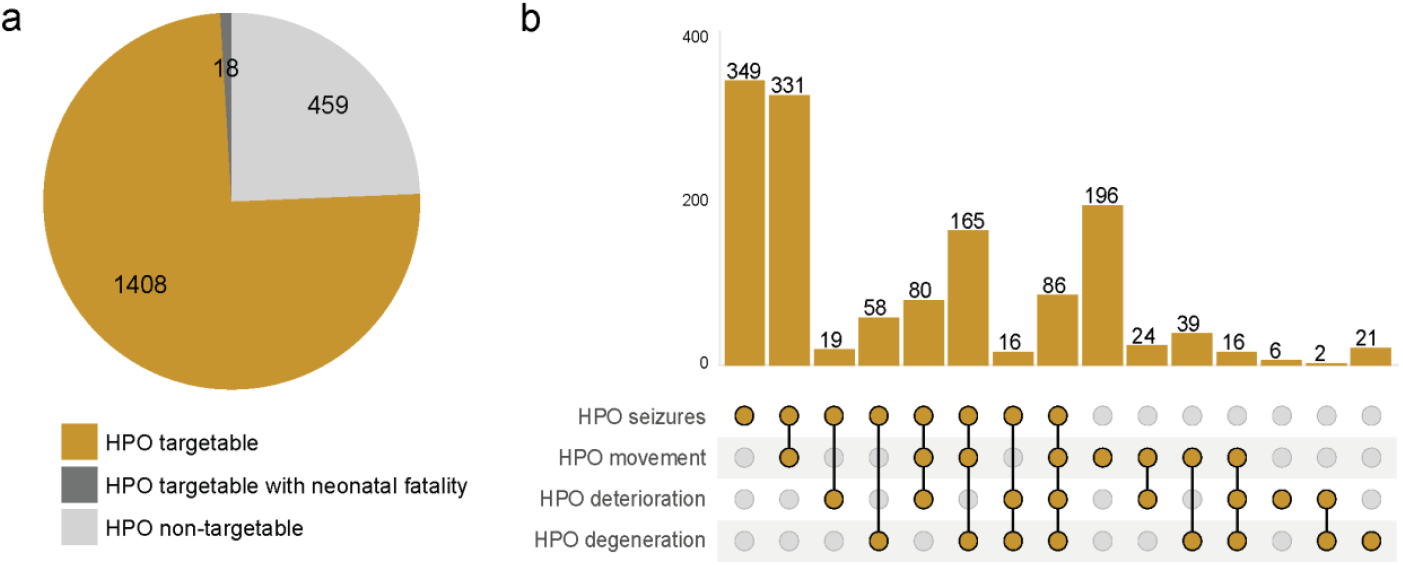
Evaluation of NDD treatability based on phenotypic features. (A) Pie-chart categorizing NDD disease-gene associations: those with at least one associated HPO term and absence of neonatal lethality (yellow), with at least one associated HPO term and neonatal fatality (dark gray), or without any associated HPO terms (light gray). (B) UpSet plot showing exclusive combinations of HPO terms associated with gene-NDD associations, where the sum across all combinations corresponds to the number of cases in the yellow section of the pie chart. HPO (Human Phenotype Ontology)

This dataset provided the basis for a bottom-up assessment of the targetability of the NDDs for different AON-based strategies. In the following section, we further explored the dataset, evaluating the presence of targetable variants, molecular and phenotypic characteristics for various AON strategies separately. All assumptions, filter criteria and variables used in the identifications of targetable NDDs and variants can be found in the supplementary data (Supplementary Table 2).

### Evaluating targetable characteristics for AON strategies

#### Deep intronic variants for splice-correction

Milasen represents a significant milestone in personalized medicine and showcases the potential of AONs to correct splice-disrupting variants, potentially more widely applicable to NDDs(3). We aimed to assess the impact of this AON strategy on NDDs using our NDD dataset. A total of 15% (9,099/60,941) of unique (likely) pathogenic ClinVar variants in our dataset are intronic and/or likely to affect splicing. However, not all splice-disrupting variants are equally suitable for AON-mediated splice correction. That is, splice-correcting AONs can correct splice-disrupting variants that cause activation of cryptic splice sites or create new splice sites, but they cannot correct splice variants that destroy a canonical splice site, whereas exon-skipping for variant exclusion and exon skipping for reading frame restoration or TANGO might. Therefore, the minimal distance between the splice-altering variant and the canonical splice site is set to be 50 base pairs (bp) away(47). Out of the 9,099 unique intronic and splice variants in our dataset, 322 variants have a VEP consequence ‘intronic_variant’ of which 217 (0.5% of all unique variants, involved in 150 NDDs) are at least 50 bp away from the nearest canonical splice site and are therefore potentially targetable for splice-correction AON therapy (Fig. 4A, Supplementary Data 1,2).

**Fig. 4.**
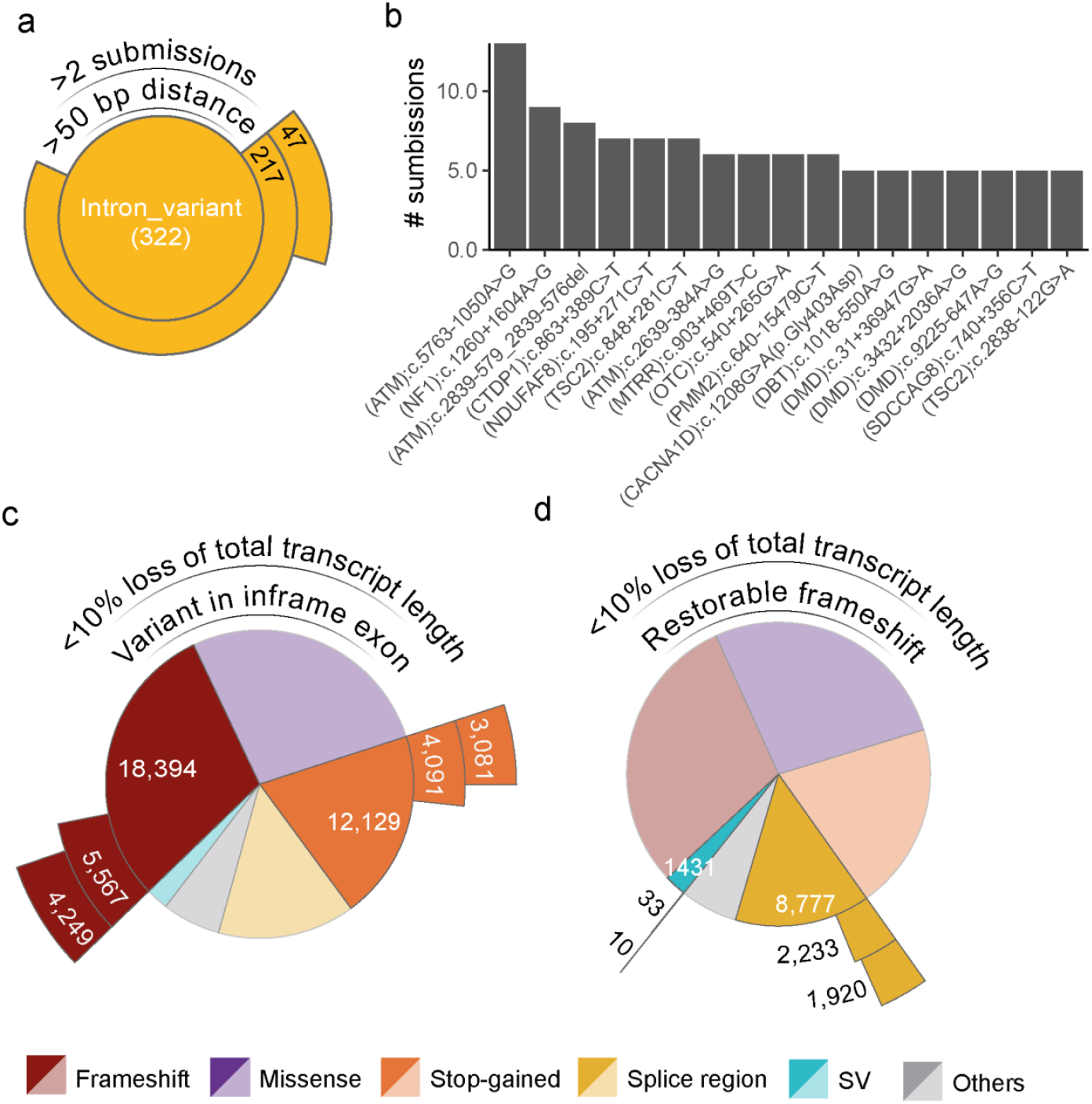
Evaluation of NDD treatability based on molecular consequences of variants. A. Sunburst plot of all variants with annotated consequence ‘intron_variant’ showcasing the applicability of splice-correcting AONs. B. The number of submissions of all ClinVar variants with 5 or more submissions. C/D Sunburst plots showcasing the applicability of exon-skipping for variant exclusion (C), and exon-skipping for reading frame restoration (D). SV: structural variant.

Developing an individualized AON therapy for individuals with NDD with a targetable splice-switching variant is challenging and expensive. Targeting recurrent splice-disrupting variants provides a more feasible approach(46). To identify recurrent targetable splice-disrupting variants, we searched our NDD dataset for splice variants with >2 ClinVar submissions and found 47 recurrent splice variants (involved in 34 NDDs) (Fig. 4A/B). These include two previously reported likely targetable deep intronic variants in ATM: NM_000051.4(ATM):c.5763-1050A>G and NM_000051.4(ATM):c.2839-579_2839-576del (Fig. 4B, Supplementary Data 1,2)(46).

#### LoF variants amenable for exon-skipping to enable shortened transcript expression

Exon skipping can be used to prevent NMD by enabling shortened transcript expression and partially restoring protein function. This can be achieved by exon skipping for variant exclusion and exon skipping for reading frame restoration (Fig. 1B,C). The suitability to use either of these two approaches for an NDD depends on the underlying variant type and is therefore discussed separately.

To evaluate to which extent exon skipping for reading frame restoration might be used for NDDs, we aimed to identify mutations that result in full exon deletions predicted to create a frameshift and which - by skipping an adjacent exon using an AON - would restore the reading frame. Additionally, we assumed that splice variants within less than three bp from a canonical splice site likely induce the skipping of the adjacent out-of-frame exon, essentially achieving the same effect as the deletion of a full exon. In our NDD dataset, structural variants (SVs) (n=1,431) and splice region variants (n=8,777) comprise 17% of the unique variants (Fig. 4D). Of the 1,431 SVs, 109 (0.2%) were whole-exon deletions leading to a frameshift. Additionally, we observed that 30% (2,611/8,777) of all splice region variants are splice donor or acceptor variants located at the canonical splice site positions (+/- 2bp) of an out-of-frame exon. Subsequently, we selected variants for which skipping an adjacent exon would restore the reading frame. Notably, skipping a biologically relevant transcript’s first or last exon is impossible, thereby we excluded splice variants affecting the first and last exons for exon skipping for reading frame restoration. These combined criteria left 33 (0.05%) full exon frame-shift SVs involved in 29 NDDs and 2,233 (3.7%) splice variants involved in 597 NDDs that might benefit from exon skipping for reading frame restoration (Fig. 4D).

Exon skipping for variant exclusion prevents the incorporation of an in-frame exon containing an NMD-triggering LoF variant to allow the expression of a shortened gene product and escape of NMD. In our NDD dataset, 30,523/60,941 (50%) of the NDD-causing variants are stop-gain variants or out-of-frame indels (Fig. 4C). Excluding those located in out-of-frame exons and the first and last exon for biological reasons, left 9,658/60,941 (16%) of the ClinVar variants, involved, in 1,128 NDDs potentially amenable for exon-skipping for variant exclusion (Fig. 4C).

As a rule of thumb, exon skipping that results in more than a 10% loss of the total coding sequence is not likely to produce a functional protein(48). Based on this, we further refined our selection by assessing the impact on the total length of the coding sequence, should exon-skipping for variant exclusion or reading frame restoration be applied. By selecting for variants that lead to a total loss of coding sequence length of 10% or less, the number of targetable variants was reduced from 2,266 to 1,930 (3% of total unique variants) in 470 NDDs for exon-skipping for reading frame restoration and from 9,658 to 7,330 (12%) in 754 NDDs for exon-skipping for variant exclusion (Fig. 4C/D; Supplementary Data 1/2). As expected, this set includes 88 ClinVar variants linked to DMD (OMIM: 310200), for which various FDA-approved exon-skipping for reading frame restoration AON therapies exist(5, 49), suggesting that our approach to select NDD variants amenable to exon skipping for reading frame restoration is feasible.

#### NDDs caused by haploinsufficiency allowing for wildtype allele upregulation strategies

TANGO and AON-mediated targeting of the UTR or regulatory lncRNAs upregulate expression by increasing expression of the wildtype allele (Fig. 1D,E-G). Therefore, these approaches are only applicable for NDDs caused by heterozygous LoF in haploinsufficient genes, although in some cases dominant missense variants that result in (partial) LoF or X-linked dominant variants in females might also be targetable. To identify NDDs amenable to AON strategies that increase expression of the wildtype allele, we selected autosomal dominant (AD) NDDs in haploinsufficient genes (probability of being loss-of-function intolerant: pLI > 0.9). Among the 531 AD NDDs examined, 372 were attributed to genes with a pLI > 0.9. A ratio of more than 20% pathogenic LoF/missense variants is suggestive for whether the gene causes disease through loss‐ or gain‐of‐function(37).

Applying this criteria identified 182 of 372 which are likely to cause disease through heterozygous LoF mechanism and potentially amenable for AON-strategies that increase expression of the wildtype allele (Fig. 5A). For these, we subsequently assessed the presence of the additional requirements for TANGO and AON-mediated targeting of the UTR or regulatory lncRNAs to upregulate expression separately.

**Fig. 5.**
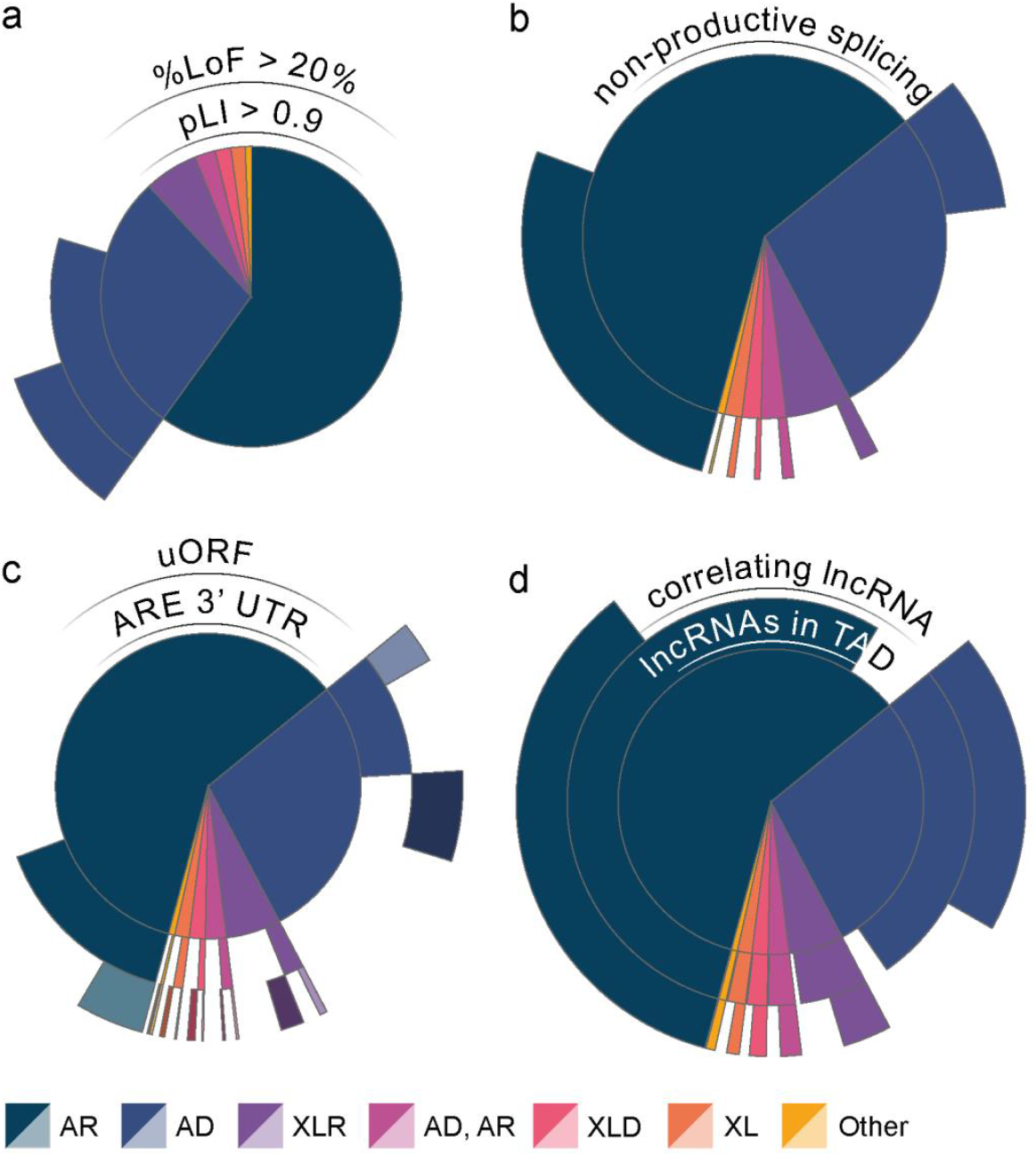
Evaluation of NDD treatability based on molecular characteristics of causative genes. Sunburst plots showcasing the number of NDDs amenable for upregulation of the wildtype allele (A), with non-productive splice products (B), regulatory structures in the 5’ and 3’ UTR (C) and the presence of correlated lncRNAs in the TAD of NDD-causing genes (D). LoF: loss-of-function, pLI: probability of being loss-of-function intolerant, uAUG: upstream open reading frame, ARE: AU-rich elements, lncRNA: long non-coding RNA, AR: autosomal recessive, AD: autosomal dominant, XLR: X-linked recessive, XL: X-linked recessive or dominant, XLD: X-linked dominant.

#### Non-productive splicing in NDD genes for the application of TANGO

The TANGO approach exploits naturally occurring non-productive splicing events to increase endogenous expression of the unaffected allele to restore protein levels in the case of haploinsufficiency(12) (Fig. 1D). To determine the possibilities for TANGO for NDDs, we intersected a set of 7,757 unique (non-)coding genes for which non-productive splice products were previously reported with our NDD dataset(12). This resulted in the identification of 724 NDDs (41%) in our NDD dataset of which the causative gene has reported non-productive splice products (Fig. 5B, Supplementary Data 1). Of these 724 NDDs, 66 NDDs were determined to be amenable for wildtype allele upregulation in alignment with the rationale employed for the identification of NDDs amenable for wildtype allele upregulation, described above (Fig. 6A).

**Fig. 6.**
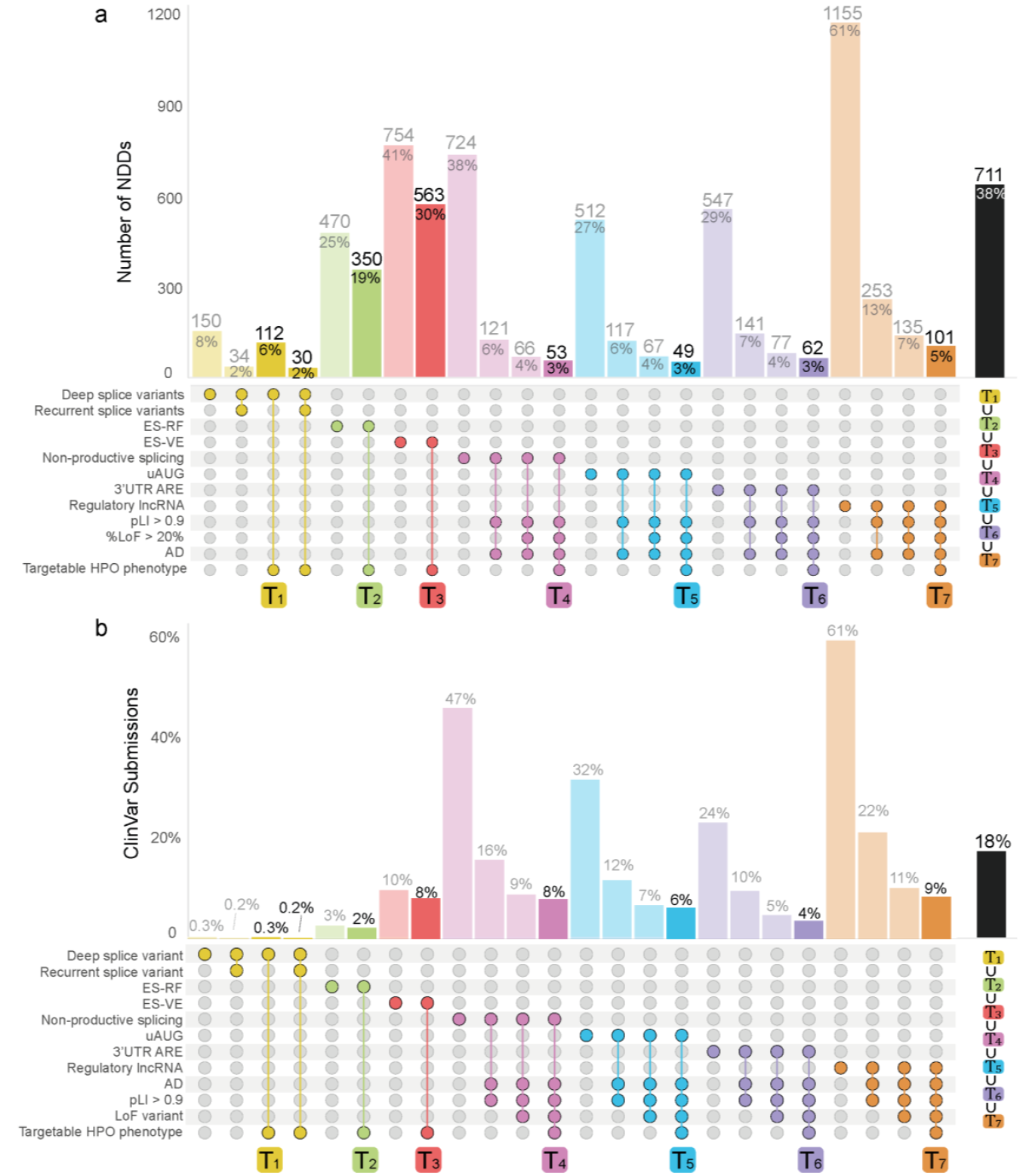
UpSet plot of favorable combinations of characteristics for AON therapy. UpSet plot of favorable characteristics on disease level (A) and patient level (represented by ClinVar submissions) (B) for seven AON therapies: Splice-correction using AONs (yellow, T_1_), exon-skipping for reading frame restoration (green, T_2_), exon-skipping for variant exclusion (red, T_3_), TANGO (pink, T_4_), uAUG targeting (blue, T_5_), 3’ ARE targeting (purple, T_6_) and AON-mediated degradation of lncRNAs (orange, T_7_). The bar representing the most favorable combination of characteristics is highlighted. The UpSet plot shows combinations of targetable characteristics associated with NDD-gene associations, where a single NDD disease-gene association can be targetable by multiple AON therapies. The union (∪), rather than the sum, of all highlighted targetable combinations (T_1-7_) (black bar) corresponds to the total number of targetable NDD-gene associations, accounting for the fact that NDD disease-gene associations can be targetable by more than one AON strategy. ES-RF: exon skipping for reading frame restoration, ES-VE: exon skipping for variant exclusion, uAUG: upstream start codon, ARE: AU- rich elements, lncRNA: long non-coding RNA, LoF: loss-of-function, pLI: probability of being loss-of-function intolerant, AD: autosomal dominant, HPO: Human phenotype ontology..

#### Regulatory elements in the UTRs of NDD genes

The role of regulatory sequences, including uAUGs and AU-rich elements, in the context of NDDs, is receiving increasing attention(10, 29). Since uAUGs and AU-rich elements involve specific sequence motifs, computational identification of uAUGs and AU-rich elements is feasible, which has led to several databases containing their locations(39, 40). To identify NDDs that are potentially amenable to a UTR-targeting AON strategy, we annotated our dataset of NDDs with these predictions(39, 40). This analysis identified 512 (27%) and 547 (29%) NDDs for which the causative gene has one or more predicted uAUGs or 3’UTR AU-rich elements, respectively (Fig. 5C, Supplementary Data 1). Selecting only the NDDs amenable for wildtype allele upregulation, as described above, identified 67 and 77 NDDs as potential targets for uAUGs or 3’UTR AU-rich element- targeting AONs respectively (Fig. 6A).

#### Regulatory cis-acting lncRNAs of NDD genes

To identify possible cis-acting lncRNAs that can be used to upregulate the expression of NDD genes, we determined for all NDD genes whether lncRNAs are located within the same topologically associated domain (TAD) as defined in TADmap(41, 42). This revealed that 1,733 (92%) of the NDDs in our dataset are caused by variants in a gene that have at least one lncRNA in their TAD (Fig. 5D). Next, we assessed whether the lncRNAs were co-expressed with the NDD genes of interest within its TAD using brain expression data cataloged in GTEX(43). We identified significant co-expression for 1,155 (58%) NDD genes and at least one lncRNA nearby (Fig. 5D, Supplementary Data 1). As expected, this set included *UBE3A* and *CHD2*, being two genes that have been upregulated by AON-mediated degradation of their regulatory cis-acting lncRNAs, *UBE3A-ATS*(50) and *CHASERR*(42), respectively. Consistent with our prior strategy, we selected only NDDs amenable to wild-type allele upregulation. This identified 135 NDDs as potential targets for targeting regulatory cis-acting lncRNAs with AONs (Fig. 6A).

### The potential impact across the spectrum of AON strategies for NDD

To determine the overall number of NDDs that are suitable for any AON therapy, we next integrated the data obtained from each of the individual analyses (Supplementary Data 1,2). Combining the relevant targetable traits for each AON therapy revealed that 38% (n = 711) of the NDDs have targetable characteristics for at least one AON therapy (Fig. 6A). However, the amenability for AON therapy can be influenced by the specific genetic variant present in each affected individual, even if the NDD itself exhibits targetable characteristics for AON therapy. To estimate the impact of each AON strategy on an individual level we took the number of ClinVar submissions for each reported variant as a measure of occurrence. This revealed that 18% (23,158/125,380) of ClinVar submissions have targetable characteristics for at least one AON strategy (Fig. 6B).

## Discussion

AON therapy for NDDs is a fast-growing field of interest uncovering considerable uncharted potential. We explored the presence of targetable phenotypic features, predicted underlying variant consequence and gene properties of 1,885 NDDs and their causative genes to estimate the impact of AON therapy for NDDs and to identify promising candidates for future AON therapies. Doing so, we identified that AON-based therapeutic strategies may be possible for an estimated 18% of the individuals across 711 (38%) different NDDs, which demonstrates that there might be a more extensive potential for the use of AON therapy in NDDs than was anticipated thus far. Among these potentially targetable NDDs and variants are AON strategies that show promising results in preclinical studies or tentative success in clinical trials including the TANGO strategy for Dravet syndrome and *SYNGAP1* syndrome (OMIM: 612621), exon skipping for reading frame restauration for *DMD* and AON-mediated degradation of lncRNAs for Angelman syndrome and *CHD2* Syndrome (OMIM: 615369). The (re)identification of these NDDs with demonstrated potential for AON-based intervention suggests that AON-based therapeutic strategies may be possible for at least a part of the individuals with the other potentially targetable NDDs for which currently no approved therapy exists. These include NDDs like Coffin-Siris syndrome (*ARID1B*; OMIM: 105830), Kleefstra syndrome (*EHMT1*; OMIM:610253) and Koolen-De Vries syndrome (*KANSL1*; OMIM: 610443). Still, it is important to note that this study is exploratory in nature, carrying inherent uncertainties that may over- or underestimationthe potential of an AON strategy and requires consideration both in the research and clinical contexts. Although we cannot yet precisely quantify the impact of AON strategies, we provide a framework for predicting their applicability. This framework can guide future research and be refined as more data are collected, improving our ability to identify NDDs and variants that may be suitable targets for AON therapy.

More extensive assessment of variant consequences, gene properties and pathological mechanisms of genetic disorders can refine the identification of targetable NDDs and variants. For instance, *in silico* prediction based on spatial clustering of variants and missense-to-loss-of-function ratio in combination with tools that can help predict the functional consequences of missense variants such as AlphaMissense hold promise to expand our bottom-up approach to identifying NDDs suitable for AON-mediated degradation of overexpressed, overactive, or toxic genes(36, 37, 51). Furthermore, deep-learning based tools like splice-AI can help identify exonic splice variants or intronic splice variants <50bp away from a splice-junction that are amenable for AON-mediated splice-correction(52). We also note that although we show that 8% of the variants can be potentially targeted by exon skipping for variant exclusion, in reality, this approach will often lead to a non-functional protein even if less than 10% of the protein length is lost. Integrating genetic conservation data and utilizing protein folding tools like AlphaFold can help predict whether protein functionality is retained, but skipping exons that encode critical domains may still render the protein non-functional. Careful assessment of these regions is therefore essential. For these and other AON strategies, advanced sequencing techniques can further refine the selection of NDDs and patients amenable to AON therapy. This includes high-throughput reverse genetic screening for lncRNAs, Riboseq for the identification of uAUGs and short- and long-read RNA sequencing to detect and quantify non-productive splice events for TANGO(53-55). Current variant databases, such as ClinVar, still heavily rely on data primarily derived from exon-based sequencing. The incremental use of genome sequencing will help to address the underestimation of patients amenable to splice correction as these will also allow the systematic identification of intronic variants amenable to AON-mediated splice correction. RNA sequencing can then help clarify the effects of splice variants, which often produce multiple aberrant transcripts, thereby guiding more precise AON-mediated splice correction. Conducting these studies in the relevant tissue and cell types is of high importance, as the presence of favorable characteristics for AON strategies can vary temporospatially, significantly impacting the efficiency and therapeutic potential of AON therapy(13, 56).

One of the main hurdles to overcome when identifying NDDs amenable for AON therapy is determining which clinical phenotype(s) can be (sufficiently) rescued. In our analysis, we considered seizures, progressive phenotypes, and movement disorders as markers of targetability, but each comes with unique limitations. For instance, seizures linked to neuroanatomical defects are unlikely to improve without addressing the underlying structural abnormalities. NDDs associated with movement disorders that stem from peripheral defects - such as those involving the neuromuscular junction - may require systemic administration or novel delivery vehicles. The assessment of rescuable clinical phenotypes is further complicated by the clinical landscape of NDDs, marked by a broad spectrum of phenotypic features and clinical heterogeneity. For example, variants in ceroid lipofuscinosis, neuronal, 10 (*CLN10;* HGNC:2529; OMIM: 610127) often lead to a congenital, and most likely, untreatable form of Batten disease resulting in neonatal lethality. However, in some cases, *CLN10*-associated Batten disease represents a potentially targetable neurodegenerative disorder with a juvenile-onset(57, 58), that is similar to the *CLN7*-type Batten disease, for which the AON therapy Milasen has been reported to show clinical benefits in a single patient(3).

The wide variability in severity of clinical features of NDDs also raises ethical considerations for AON therapy. Currently, AON therapies target severely debilitating or fatal NDDs, which was not considered into our phenotypic selection criteria. Other considerations should include whether a condition’s severity justifies therapy, especially when AON therapy may not extend to mitigating other, sometimes more severe, clinical (co)morbidities of the condition. In some of these instances, however, the development of comorbidities may be prevented, suggesting that, for instance, starting preventative therapy as early as possible would be most beneficial. Examples of these are presented such as the early treatment of SMA upon identification in newborn screening(59, 60).

Taken together, the selection of NDDs with a targetable phenotype is and will remain complicated. Eventually, the decision to pursue the development and administration of AON therapy should be a collaborative effort between parents and physicians, carefully considering the balance between risks and potential benefits for each NDD and possibly each patient. Extensive in vitro and in vivo studies have to be performed for each potentially targetable NDD, and in some cases each individual, to confirm the therapeutic potential of each AON strategy. On a functional level, personalized drug screening with (patient-derived) iPSC-derived models combined with functional studies can be utilized. However, due to the often, complex biology and poorly understood etiology of NDDs, this process will be lengthy and complicated, especially for less-prevalent and less-studied NDDs.

Moreover, scaling these approaches remains a significant challenge, given the resources and time required for each individual case. In light of these challenges, we estimated that gene-based AON strategies, like TANGO or targeting of the UTR/regulatory lncRNAs, can potentially be used for 9-60 times more patients using one AON compared to variant-based AON strategies, like splice-correction and exons skipping for variant exclusion and frameshift restoration (Supplementary Table 1). Hence, the use of gene-based AON strategies requires fewer resources and expenses to treat an equivalent number of individuals. It is, however, also important to note that the impact of variant-based AON strategy highly depends on the selected variant, as was also indicated in a previous study(46).

Focusing on only recurrent variants, splice-correcting AONs can potentially treat 2.5x more patients compared to a non-focused approach, making the development of splice-correcting AONs more feasible (Supplementary Table 1).

Nevertheless, the limited applicability of each AON, especially for ultra-rare NDDs and variant-based AON strategies, poses a significant challenge. Clinical trials are required to validate safety and efficacy in small and diverse patient populations, or even, single patients. Particularly for these N=1/few trials, standardized evaluation methods and outcome measures are still lacking, challenging their therapeutic assessment. This is further complicated by the limited understanding of the natural history of many NDDs and the lack of reliable outcome measures. For instance, seizures can improve spontaneously, potentially confounding their use as a reliable outcome measure in clinical trials. In line with the goal of the International Rare Diseases Research Consortium (IRDiRC, https://irdirc.org/) to develop 1,000 therapies for (ultra) rare diseases and/or n-of-1 cases, collaborative efforts, such as the N=1 Collaborative (https://www.n1collaborative.org/), the 1 mutation 1 medicine (1M1M, https://www.1mutation1medicine.eu/) collaborative, the collaborative Dutch Center for RNA Therapeutics (DCRT, https://www.rnatherapy.nl/) and the nLorem Foundation (https://www.nlorem.org), play a pivotal role in supporting and enabling AON therapy(61, 62).

Overall, our study presents a framework for evaluating AON strategies for monogenic NDD therapy, adaptable to future insights, and suggests a broader applicability of AON therapy beyond the select group of NDDs and individuals for which AON therapy is currently applied. The methodology we employed to assess the amenability of different AON strategies can be extended to identify other categories of monogenic disorders suitable for AON therapy, such as inherited eye-related disorders, contingent upon adapting the definition of targetable phenotypic presentations. While our findings underscore the promise of AON therapies, it is important to recognize that the field is still in its early stages, with relatively few examples of proven clinical efficacy. Further studies, can and should, help to refine the identification of targetable disorders and variants to confirm or disprove their targetability, both on a functional, as well as on clinical level. Eventually, these collective efforts can provide us with the necessary tools to uncover the true potential of AON therapies for monogenic disorders and advance their development, potentially improving the quality of life for many affected individuals with NDDs and their families.

## Conclusions

We found that AON-based therapeutic strategies could be feasible for an estimated 18% of individuals across 711 (38%) different NDDs. Among these are AON strategies for NDDs that show promising results in preclinical studies or tentative success in clinical trials. The exploratory nature of this study necessitates careful consideration in both research and clinical contexts, yet, our findings identify potentially targetable NDDs, including both well- and less-studied NDDs. Although further studies are required to validate our findings, these at minimum suggest a broader applicability of AON therapy beyond the currently targeted NDDs and individuals. Additionally, the methodology we used can be adapted to identify other categories of monogenic disorders.

## List of Abbreviations

AD: Autosomal dominant
AONs: Antisense oligonucleotides
AR: Autosomal recessive
ARE: AU-rich elements
AS: Angelman Syndrome
AT: Ataxia-telangiectasia
bp: Basepair(s)
DMD: Duchenne muscular dystrophy
DS: Dravet Syndrome
lncRNA: Long non-coding RNA
LoF: Loss-of-function
NDDs: Neurodevelopmental disorders
NMD: nonsense mediated decay
pLI: probability of being loss-of-function intolerant
PTC: premature termination codon
SMA: Spinal Muscular Dystrophy
SVs: Structural variants
TAD: topologically associated domain
TANGO: Targeted augmentation of nuclear gene output
uAUG: upstream start codon
UTR: Untranslated region
VEP: Variant effect predictor

## Ethics approval and consent to participate

Not applicable

## Consent for publication

Not applicable

## Availability of data and material

All data generated or analysed during this study are included in this published article [and its supplementary information files].

## Code availability

Datasets from different sources were integrated and deduplicated using a combination of code and manual curation. Thus, the standalone code is not sufficient to regenerate created datasets.

Therefore, this code is not available upon request. The code used for the evaluation of treatability will be made available on GitHub (URL) upon publication. Analyses were executed in R (v4.2.0).

## Acknowledgments

We thank Elke de Boer and ERN-ITHACA for discussions on the NDD gene dataset used for the evaluation of NDD therapies.

## Author Contributions

Conceptualization: K.N.W., L.E.L.M.V., N.N.K.; Writing-original draft, Data curation, Formal analysis: K.N.W.; Supervision, Writing-review & editing: L.E.L.M.V., N.N.K.

## Material & Correspondence

Corresponding author: LELM Vissers

## Supplementary Files

### Supplementary Information

**Supplementary Data 1:** Annotated NDD dataset with predicted suitable AON strategies

**Supplementary Data 2:** Annotated dataset of ClinVar variants linked to the NDD dataset with predicted suitable AON strategies

**Supplementary Data 3**. HPO terms included for targetable phenotypic feature identification

**Supplementary Note**. Detailed explanations and background information of all AON-strategies depicted in Figure 1.

