## Supplementary Information for "Systematic analysis of genetic and phenotypic characteristics reveals antisense oligonucleotide therapy potential for one-third of neurodevelopmental disorders"

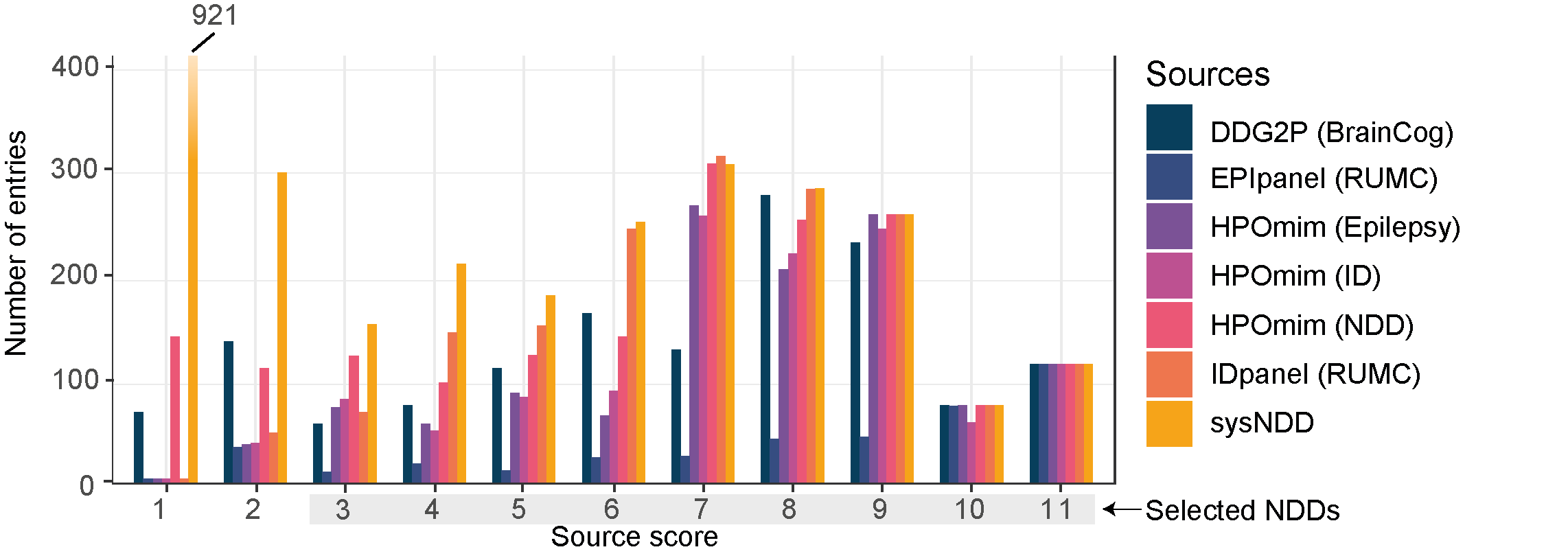

**Supplementary Figure 1.** Source score contribution per source database.

| **AON strategy** | **Number of submissions variants** | **Predicted number of AONs** | **Individuals per AON** |
| --- | --- | --- | --- |
| **Splice-correction** | 359 | 170 | 2 |
| **Splice-correction of recurrent variants** | 198 | 43 | 5 |
| **ES-RF** | 2744 | 810 | 3 |
| **ES-VE** | 10262 | 1738 | 6 |
| **TANGO** | 10499 | 91 | 115 |
| **uAUG interference** | 8305 | 69 | 120 |
| **ARE inhibition** | 4884 | 88 | 56 |
| **lncRNA targeting** | 11299 | 165 | 68 |

**Supplementary Table 1.** Average number of amenable individuals per AONs per AON strategy

| Strategy | Assumption* | Filter criteria/variable |
| --- | --- | --- |
| Splice correction | Splice-correction is only feasable intronic splice-variants that do not affect canonical splice sites can be targeted with splice-correction AONs | VEP-consequence = Intronic variants, location = >50bp away from canonical splice sites (in the MANE transcript) |
| ES-RF | ES-RF is only feasible for fully deleted exons (leading to a frame shift) or splice-variants leading to skipping of an out-of-frame exon | Variant type = deletion (of an out-of-frame exon) or VEP-consequence = splice-donor/acceptor variants (flanking an out-of-frame exon) |
|  | ES-RF is only achievable if skipping a single exon -immediately adjacent to the deletion or skipped out-of-frame exon - corrects the predicted frame shift caused by the mutation | Frame-shift restoration possible by skipping at least one adjacent exons |
|  | ES-RF does not lead to a functional protein if the first or last exon is lost by either the mutation or due to the AON | Exon (deleted or skipped by the AON) must not be the first or last exon (in the MANE transcript) |
|  | ES-RF does not lead to a function proten if more than 10% of the protein is lost. | ≤10% protein loss when mutations and skipped exon are considered. |
| ES-VE | ES-VE is only feasable if it does not affect the reading frame | Variant in a in-frame exon |
|  | ES-VE does not lead to a functional protein if the first or last exon is skipped by the AON | Exon (skipped by the AON) must not be the first or last exon (in the MANE transcript) |
|  | ES-VE is only applicable for stop-gained and frame-shift variants and not missense variants | VEP-consequence = splice-gained variant or frame-shift variant |
|  | ES-VE does not lead to a function proten if more than 10% of the protein is lost. | ≤10% protein loss when mutations and skipped exon are considered. |
| TANGO | Any detectable non-productive splice event is potentially targetable for TANGO | 1 or more non-productive splice event in the affected gene |
|  | TANGO requires the variant to lead to loss-of-function | pLI > 0.9 & >20% LoF variants (VEP-consequence = stop-gained, frame-shift or splice-donor/acceptor variant) |
|  | TANGO requires the presence of one intact allele to be upregulated | inheritance = AD |
| uORF-targeting | Any detectable non-productive splice event is potentially targetable for uORF-targeting | 1 or more uORF in the affected gene |
|  | uORF-targeting requires the variant to lead to loss-of-function | pLI > 0.9 & >20% LoF variants (VEP-consequence = stop-gained, frame-shift or splice-donor/acceptor variant) |
|  | uORF-targeting requires the presence of one intact allele to be upregulated | inheritance = AD |
| ARE-targeting | Any detectable non-productive splice event is potentially targetable for ARE-targeting | 1 or more ARE in the affected gene |
|  | ARE-targeting requires the variant to lead to loss-of-function | pLI > 0.9 & >20% LoF variants (VEP-consequence = stop-gained, frame-shift or splice-donor/acceptor variant) |
|  | ARE-targeting requires the presence of one intact allele to be upregulated | inheritance = AD |
| lncRNA-targeting | A lncRNA should regulate gene-expression of the affected allele | 1 or more lncRNA within the same TAD as the affected gene |
|  |  | Correlation in expression between lncRNA and the affected gene in GTEX brain data |
|  | lncRNA-targeting requires the variant to lead to loss-of-function | pLI > 0.9 & >20% LoF variants (VEP-consequence = stop-gained, frame-shift or splice-donor/acceptor variant) |
|  | lncRNA-targeting requires the presence of one intact allele to be upregulated | inheritance = AD |
| all strategies | AON therapy is viable for patients if the phenotypic feature can be reversed or if the therapy has the potential to halt disease progression | Disorders link to HPO-terms (in OMIM): developmental, mental or ambulation deterioration (HP:0002376, HP:0001268, HP:0002505), neurodegeneration (HP:0007367), or seizures (HP:0001250) or abnormality of movement (HP:0100022) or any HPO-term downstream of these HPO terms |
|  | The time-window for AON therapy is too short to treat NDDs causing neonatal death | Disorders link to HPO-terms (in OMIM): neonatal death (HP:0003811) are excluded |

**Supplementary Table 2. Assumptions and variables per AON strategy.** AON: Antisense oligonucleotide, VEP: Variant Effect Preditor, ES-RF: Exon skipping for reading frame restauration , ES-VE: Exon skipping for variant exclusion, TANGO: Targeted Augmentation of Nuclear Gene Output, LoF: Loss of Function, AD: Autosomal Dominant, ARE: AU-rich element, lncRNA: long non-coding RNA, TAD: Topologically associated domain, HPO: Human Phenotype Ontology. **The assumptions presented in this table were made to facilitate analyses while realizing that true complexity of both therapeutic opportunities and/or phenotypic presentation and/or biological impact is underestimated by these assumptions. They should, as such, be interpreted as approximations based on the available data, tools and information and context. With increasing insights, the analyses can be repeated in our framework by adapting the underlying assumptions and filter criteria.*
