## Supplementary Note for "Systematic analysis of genetic and phenotypic characteristics reveals antisense oligonucleotide therapy potential for one-third of neurodevelopmental disorders"

### Supplementary Note: Background information Figure 1

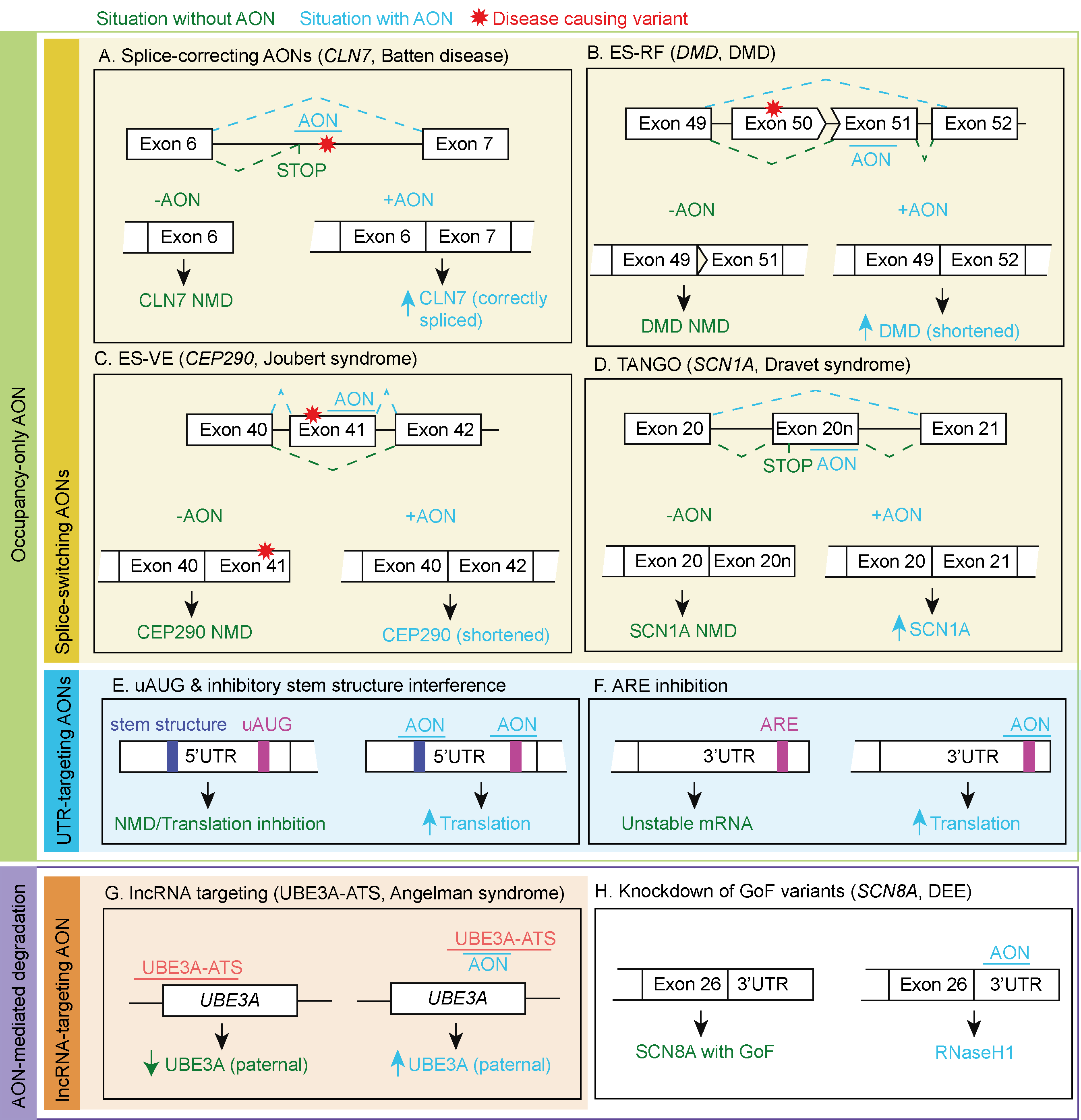

**Fig. 1. Mechanisms of AON strategies in neurological disorders.** A. Batten’s disease, the AON rescues aberrant splicing of intron 6 by steric hindrance of a cryptic splice-acceptor site(1). B. Duchenne muscular dystrophy (DMD), the AON modulates exon skipping of exon 51 directly upstream of a deleted exon 50 for reading frame restoration(2). C. Joubert syndrome, the AON mediates skipping of an exon harboring a truncating variant(3). D. Dravet syndrome, the AON prevents the production of the non-productive splice product containing the poison exon 20N and thereby upregulates expression(4). E. The AON blocks stem structures or uAUGs in the 5’UTR to increase translation efficiency(5). F. the AON blocks an ARE in the 3’UTR to increase translation efficiency(6). G. Angelman syndrome, the AON blocks the transcription of the lncRNA UBE3A-ATS and increases paternal *UBE3A* by inhibiting transcriptional interference(7). H. *SCN8A* DEE, the AON causes RNaseH1-mediated degradation of *SCN8A* mRNA(8). NMD: Nonsense-mediated decay, ARE: AU-rich elements, ES-VE: exon skipping for variant exclusion, ES-RF: exon skipping for reading frame restoration, uAUG: upstream start codon.

### Background information Figure 1

AON therapy is a highly versatile group of therapies, which can be divided into two main groups based on their mechanism of action: AONs that mediate degradation and those that occupy mRNA, creating steric hindrance (occupancy-only AONs) (FIG. 1). AON-mediated degradation selectively degrades RNA by activating RNase H1, whereas occupancy-only AONs are RNase-H1 resistant and form a steric block inferring with mRNA-protein or mRNA-oligonucleotide binding (9). Occupancy-only AONs are primarily used to alter pre-mRNA splicing (splice-switching AONs) but can also be used to alter the RNA structure or prevent the binding of regulatory proteins in coding regions and untranslated regions (UTR) (1, 4, 5, 7, 10)(FIG. 1). In this supplementary note, we will discuss the mechanisms of current AON strategies and how they have been employed in treating neurological disorders.

#### The use of splice-switching AONs

Splice-switching AONs prevent recognition of an authentic or cryptic splice site by binding to the authentic or cryptic splice-acceptor or -donor site, exonic or intronic splice enhancers (ESEs/ ISEs), or exonic or intronic splice silencers (ESSs/ISSs). Nusinersen, a splice-switching AON used to treat SMA, was the first AON-based therapy to treat a rare genetic disease of the CNS by direct administration to the CNS. SMA is caused by autosomal recessive loss-of-function (LoF) variants in the *SMN1* gene (HGNC:11117). *SMN2* (HGNC:11118) is nearly identical to the *SMN1* gene (by percentage sequence homology and function), but only a small fraction of *SMN2* leads to functional protein due to a single nucleotide variant (SNV) in intron 7. This SNV causes skipping of exon 7, leading to nonsense-mediated mRNA decay (NMD) of the *SMN2* gene product. Nusinersen binds to an ISS in the *SMN2* pre-mRNA and alters the splicing of *SMN2* pre-mRNA to include exon 7, enabling the translated SMN2 protein to replace the function of missing SMN1 protein in individuals with SMA (11). Treatment of Nusinersen halted disease progression and improved motor function of individuals with SMA during clinical trials (12). Hence, in 2016, the FDA prematurely approved Nusinersen after positive interim results for individuals with SMA. The success of Nusinersen boosted the use of AONs to treat genetic diseases and marked the starting signal for the use of splice-switching AONs to treat neurological diseases. Currently, splice-switching AONs are employed to treat neurological diseases by rescuing splice-disrupting variants, exon skipping to restore the gene function of a variant-containing allele and to increase levels of wild-type mRNA transcripts to boost functional protein levels (1, 2, 4).

##### Rescuing splice-disrupting variants with splice-switching AONs

Splice-disrupting variants can cause various splicing abnormalities, such as exon skipping, intron retention, activation of cryptic splice sites, or creation of new splice sites. To counteract these variants, splice-switching AONs aim to restore the disrupted splicing. For instance, in an individual with CLN7-type Batten disease, an SNV in intron 6 created a cryptic splice-acceptor site in *MFSD8* (HGNC:28486), leading to the inclusion of a pseudo-exon leading to NMD (1). An AON called Milasen, which targets the introduced splice acceptor site, increased the relative abundance of the canonical splice product and partially restored the MFSD8’s vacuole accumulation *in vitro* (FIG. 1A)(1). Although not all of the phenotype could be reversed, a clinical trial showed that Milasen reduced the seizure frequency and length by 50% and stabilized global motor function and sensory threshold testing upon administration to the person (1).

Rescuing splice-disrupting variants using AONs is not limited to Milasen. Another example is the rescue of altered splicing by an exonic variant in exon 55 of *ATM* (NM_000051.4(ATM):c.7865C>T, HGNC:795) by a splice-switching AON, Atipeksen, used in individuals with ataxia-telangiectasia (AT, OMIM: 208900) (13). Although this AON successfully rescued disrupted splicing of the *ATM* transcript, whether this results in clinical improvements is currently unknown. Unlike the variant treated with Milasen, this variant was found to be recurrent in individuals with AT and may therefore be used for multiple individuals with AT harboring the c.7865C>T variant (14, 15). The tentative success of Milasen and Atipeksen shows that using individualized AONs to treat neurological disorders is feasible and might be used for individuals with a neurological disease caused by a splice-disrupting variant.

##### Exon skipping to restore gene function of a variant-containing allele

Splice-switching AONs can also be used to skip exons to restore the reading frame, or alternatively, they can be used to skip an exon harboring a pathogenic variant (FIG. 1B/C). To distinguish the two methods, we termed these AON strategies exon skipping for reading frame restoration (ES-RF) and exon skipping for variant exclusion (ES-VE), respectively.

ES-RF involves the skipping of an out-of-frame exon located in the exon upstream or downstream of the pathogenic variant, causing a reading frame shift to restore the reading frame and produce a shorter protein in which the amino acids encoded by the skipped exon(s) are lacking. This AON strategy has demonstrated promising results for DMD(2). Hemizygous pathogenic variants in *DMD* (HGNC:2928) leading to a null-allele cause DMD. However, if a pathogenic variant maintains the open reading frame, the milder phenotype of Becker muscular dystrophy (OMIM: 300376) is observed. Etiplirsen is a systemically administered splice-switching AON that induces Exon 51 skipping in individuals with DMD, amenable with pathogenic variants directly upstream or downstream of exon 51, restoring the reading frame of the *DMD transcript* (FIG. 1B)(2). Eteplirsen partially restores DMD protein function, transitioning the phenotype to be similar to Becker muscular dystrophy. During a clinical trial, individuals with DMD treated with Eteplirsen displayed a significant increase in the percentage of dystrophin-positive fibers and little to no progression to loss of ambulation (2). Following this, Eteplirsen was approved by the FDA(16). However, market authorization was rejected by its European counterpart, EMA, due to a lack of clinical improvement (17). Nonetheless, clinical trials for nine drugs, targeting DMD exon 44, 45, 51, or 53, have since been proposed, of which four have reached FDA approval(16). This includes Golodirsen and Viltolarsen, which both demonstrated clinical benefit. The rising numbers of studies and clinical trials on ES-RF approaches demonstrate the great potential for neurological diseases.

Alternatively, ES-VE can be used to remove an in-frame exon containing a frame-shift or nonsense variant that otherwise causes the transcript to be targeted by NMD, allowing the expression of a shorter gene product now escaping NMD (FIG. 1C). Whereas this approach has not yet been used in neurological disease, ES-VE has been shown to *in vitro* restore the cellular function of *DYSF* (HGNC:3097) associated with Miyoshi myopathy (OMIM: 254130) and *CEP290* (HGNC:29021) associated Joubert syndrome (OMIM: 610188) (3, 18) (FIG. 1C).

##### Targeted augmentation of nuclear gene output (TANGO)

Splice-switching AONs can also be used to upregulate gene expression by preventing the production of naturally occurring non-productive splice products subjected to NMD (FIG. 1D), referred to as Targeted Augmentation of Nuclear Gene Output (TANGO) (4). TANGO can be used to treat neurological diseases caused by haploinsufficiency (HI) by upregulating the expression of the functional transcript of the wild-type allele. Dravet syndrome (DS; OMIM: 607208) is caused by *de novo* variants in *SCN1A* (HGNC:10585), resulting in haploinsufficiency of the voltage-gated sodium channel subunit NaV1.1 channel (19). A naturally occurring poison exon (20N) is included in 89% of the wild-type *SCN1A* transcripts, resulting in a frame-shift and subsequent NMD (4). In DS mice, an AON targeted to prevent the inclusion of this poison exon was able to increase the productive transcript expression of the wild-type allele, reducing the seizure incidence and sudden unexpected death (FIG. 1D). Interim results of clinical trials in individuals treated with this AON also showed reduced seizure frequency.

Interestingly, this approach is not limited to *SCN1A*/Dravet Syndrome, as recently, a non-productive transcript of *SYNGAP1* (SYNGAP1 Syndrome; OMIM: 612621, HGNC:11497) was identified (20). Here, an alternative 3’ splice site of exon 11 leads to NMD. *In vitro*, using iPSC-derived cerebral organoids, an AON has been shown to inhibit the non-productive event and upregulate SYNGAP1 protein levels. Future clinical trials must reveal the therapeutic potential in individuals with SYNGAP1 syndrome, but the overall promise for TANGO is immense, as it can be used for all diseases caused by HI, provided that a naturally occurring non-productive splice event exists.

#### UTR-targeting AONs

UTRs, both at the 5’and 3’-end, are rich in regulatory elements, such as upstream start codons, stem structures, AU-rich elements (AREs) and binding sites of non-coding RNA, of which each element can be targeted by occupancy-only AONs to upregulate mRNA translation efficacy.

At the 5’UTR, alternative start codons (uAUGs) create upstream open reading frames (uORFs). The translation of these uORFs can inhibit translation efficiency by competing for the main translation initiation site, and if the uORF creates an out-of-frame product, it will trigger NMD (21). AONs targeting the uAUGs have been shown to increase translation efficiency by 30-150% for four genes, albeit these genes are unrelated to neurological disease (FIG. 1E) (22). Like uAUGs, tertiary stem structures in the 5'UTR can also affect the mRNA accessibility for the translation machinery and thereby inhibit translation. Targeting stem-structured inhibitory elements of three genes unrelated to neurological development by AONs, upregulated protein levels up to 2.7-fold (FIG. 1E) (5).

For the 3’UTR, alternative pre-mRNA polyadenylation signals, AREs, can contain destabilization motifs responsible for the rapid degradation of the mRNA. Targeting these destabilizing motifs by AONs can increase mRNA stability and protein expression (FIG. 1F)(6). For *SHANK3* (HGNC:14294), a gene that causes the neurodevelopmental disorder Phelan-McDermid Syndrome (PMDS, OMIM: 606232), successes have been achieved by modulating the 3’UTR. Using iPSCs-derived motoneurons from individuals with PMDS and AONs targeting the 3’UTR, SHANK3 protein levels could increase 1.3 to 1.6-fold *in vitro*, although the exact mechanism underlying this increase is unknown (10). Altogether, there seem to be good opportunities for modulating 5’ and 3’ UTRs to upregulate gene expression for disorders caused by haploinsufficiency (HI). However, so far this has not been achieved in genes related to neurological diseases and the extent to which modulating 5’ and 3’ UTRs can provide treatment options for neurological diseases remains to be determined by functional *in vitro* or *in vivo* assessment of models for neurological diseases.

#### AON-mediated degradation by RNase H1

In addition to various forms of occupancy-only AONs, AONs that mediate degradation are also gaining attention, especially for diseases caused by overexpression, overactivity or toxicity of the affected gene or downstream consequences. In essence, AON-mediated degradation aims to restore the distorted biological balance by directly targeting the affected gene/protein product or indirectly by interfering with regulators of the pathway in which the affected gene/protein product is involved, such as via lncRNAs.

##### AON-mediated degradation of overexpressed, overactive or toxic gene products

SNVs or structural variants, such as gene duplications, can induce overexpression, overactivity or lead to acquired toxicity in the function of a protein or mRNA, causing disease. For instance, gain-of-function (GoF) variants in *SCN8A*, *SCN2A*, and *KCNT1* (HGNC:10596, 10588, 18865) can lead to developmental and epileptic encephalopathies (DEE; OMIM: 614558, 613721, 614959, respectively). For these DEEs, the GoF variants lengthen the time ion channels are open, creating increased neuronal activity due to the constant flow of sodium ions into the neurons, followed by seizures. In mice with DEE, AON-mediated mRNA degradation of *SCN8A*, *SCN2A*, or *KCNT1* mRNA, respectively, were indeed able to reduce mRNA levels of the respective genes and protein levels, *in vitro* and *in vivo* (FIG. 1H)(8, 23, 24). Administration of these AONs reduced the number of seizures and extended the survival of the DEE mice models. This approach has, amongst others, also been successfully applied in rodent animal models for Schuurs-Hoeijmakers syndrome (*PACS1*, OMIM: 615009), Alexander disease (*GFAP*, OMIM: 203450), Myotonic dystrophy 1 (*DM1*, OMIM: 160900) and X-linked, Lubs-type Intellectual developmental disorder (*MECP2* duplication; OMIM: 300260)(25-28). To date, clinical trials have been started for one AON targeting *SCN2A* mRNA to treat DEE (ClinicalTrials.gov ID : NCT05737784), one targeting *GFAP* (HGNC:4235) mRNA for Alexander disease (ClinicalTrials.gov ID : NCT04849741) and clinical trials are ongoing for DM1 (ClinicalTrials.gov ID : NCT05481879). Of note, the AONs used in these examples target both the wild-type and mutant allele, resulting in the degradation of both mutated and functional gene products. This non-specificity is a cause for concern, as the combined knockdown of mutant and wild-type alleles may lead to a net haploinsufficiency or mimic null-alleles, which may result in an allelic disorder caused by HI(29). AONs that target the mutated allele may overcome this side effect. In the case of Machado-Joseph disease (OMIM: 109150), caused by variants in *ATXN3* (HGNC:7106)*,* a promising approach involves an allele-specific AON that targets a common single nucleotide polymorphism located on the mutant allele. This strategy effectively reduced the expression of the mutant ataxin-3 by 80% while leaving the wild-type ataxin-3 unaffected (30), showing future potential for other allele-specific AON-mediated degradation therapy.

##### AON-mediated degradation to restore disease pathway or system imbalance

Alternatively, AON-mediated degradation can target disease pathways or systems to compensate for dysfunctional genes related to the same pathways or systems(31). This approach is currently used for DEEs(31). An imbalance of inhibitory and excitatory neuron activity causes DEEs. As mentioned before, GoF variants in *SCN8A* cause DEE. The imbalance of inhibitory and excitatory neuron activity is due to hyperactivity in excitatory neurons carrying *SCN8A* GoF variants (8). However, LoF variants in *SCN1A*, *KCNQ2*, and *KCNA1* (HGNC:10585, 6296, 6218) can also cause DEE by reducing the firing of inhibitory neurons, leading to DEE (OMIM: 607208, [613720](https://www.omim.org/entry/613720), 160120, respectively). Previously, we mentioned that AON-mediated degradation of *SCN8A* in rodent models carrying *SCN8A* GoF improved seizure occurrence. Alternatively, inducing degradation of *SCN8A* mRNA with the same *SCN8A*-targeting AON also protected against seizures and increased survival in mice carrying LoF variants in *SCN1A*, *KCNQ2* and *KCNA1* (8, 31). These studies demonstrate that AONs can also be used to target disease pathways or system balance in DEE and possibly other neurological diseases.

##### AON-mediated degradation of expression regulatory factors

So far, the studies we have described have been focused on the down- and upregulation of elements directly targeting the mRNA of protein-coding genes. However, AONs can also target elements involved in mediating the expression levels of protein-coding genes. One such group is non-coding RNAs, specifically the cis-acting long non-coding RNAs (lncRNAs), typically representing transcripts of 200 nucleotides or more in size that are not translated into protein (32). Cis-acting lncRNAs regulate gene expression of neighboring genes at their site of transcription and AON-mediated degradation of repressive cis-acting lncRNAs can, for instance, upregulate the expression of a wild-type allele in the case of HI (FIG. 1G). This principle has been successfully demonstrated for the cis-acting lncRNA *CHASERR* (HGNC:48626), which negatively regulates *CHD2* (HGNC:1917) expression. HI of *CHD2* causes DEE type 94 (OMIM: 615369). AON-mediated degradation of the *Chaserr* lncRNA in a mouse cell line has been shown to upregulate *Chd2* mRNA expression and Chd2 protein levels (33), making this a potential treatment for individuals with DEE type 94. Another example is AON-mediated degradation of the cis-acting lncRNA, *UBE3A* antisense transcript (*UBE3A-ATS,* HGNC:37462), for Angelman syndrome (AS; OMIM: 105830), which is caused by functional loss of the maternal copy of the imprinted *UBE3A* gene (HGNC:12496). Under normal physiological circumstances, the paternal *UBE3A* gene is silenced by the *UBE3A-ATS*. Targeting this *UBE3A-ATS* lncRNA with AON-mediated degradation in a mouse model of AS undid paternal *UBE3A* silencing, thus leading to *UBE3A* expression. In mice treated with this AON, a clinical reversal of the memory impairment and the obesity phenotype were observed (FIG. 1G) (34). Currently, several clinical trials are ongoing for AS in which AONs mediated the degradation of *UBE3A-ATS* (ClinicalTrials.gov ID : NCT05127226, NCT04259281, NCT05531890) and interim data show that in individuals with AS, *UBE3A* expression was increased and clinical parameters improved.

Whereas these clinical trial results are promising, knock-out of lncRNAs *UBE3A-ATS* for AS and *CHASERR* for DEE 94 have recently shown that the underlying biological mechanisms are more complex than initially thought(7). Whereas both CHD2 and UBE3A, RNA and protein levels increased after deletions of *CHASERR* or *UBE3A-ATS*, reintroduction of the lncRNAs, *CHASERR* and *UBE3A-ATS,* could not undo this increase in *UBE3A* and *CHD2* expression(7). Further studies showed that AON-mediated degradation of *UBE3A-ATS* caused the disengagement of RNA Pol II from upstream of *UBE3A-ATS* and increased RNA Pol II transcribing of *UBE3A* (7), critical for *UBE3A* upregulation upon *UBE3A-ATS* AON-mediated degradation. These results indicate that AON-mediated degradation of cis-acting lncRNAs acts through transcriptional termination and fundamentally differs from AON-mediated degradation strategies directly targeting the affected genes.
